# Salient images: Evidence for a component process architecture for visual imagination

**DOI:** 10.1101/739953

**Authors:** Charlotte Murphy, Nerissa Siu Ping Ho, Boris Bernhardt, C. Neil Macrae, Mladen Sormaz, Deniz Vatansever, Elizabeth Jefferies, Jonathan Smallwood

**Author notes:** Corresponding Author: Charlotte Murphy, PhD, Department of Psychology, University of York, York, UK, YO105DD.

## Abstract

The ability to visualise different people and places in imagination is a core element of human cognition. Contemporary neuroimaging research highlights regions of primary visual cortex as important in instantiating images in the mind’s eye. Here we combine task-based functional magnetic resonance imaging with measures of intrinsic brain organisation to show the ventral attention, or salience, network, also plays an important role in visual imagination. In a task-based study we replicated prior observations of regions of medial occipital cortex, including Brodmann Area 17, showing greater neural activity during imagination than when perceiving similar stimuli. In addition, we found regions of the ventral attention network, and in particular right dorsolateral prefrontal cortex (BA 9/46), were activated by the same acts of imagination. In a subsequent resting-state study, we demonstrated medial occipital cortex regions form a functional network with regions of the ventral attention network that were also active during imagination. Furthermore, this pattern was maximised in an overlapping region of right dorsolateral prefrontal cortex for individuals whose self-reported imagery had stronger negative affect. Together these data suggest visual imagination relies on functional interactions between regions of primary visual cortex with the ventral attention system, and in particular, with a region of right dorsolateral prefrontal cortex.

## 1. Introduction

Human cognition is not limited in scope by immediate sensory input; imagination allows experience to encompass people, places and objects that are absent from the here and now. Contemporary psychological accounts of visual imagery, suggest that at least parts of the process through which we create an image in our minds’ eye depends upon processes that are also important in acts of perception. Neuroimaging has revealed activation within primary visual cortex during visual imagery (Chen et al., 1998; Klein et al., 2000; Kosslyn et al., 1995), while application of Transcranial Magnetic Stimulation (TMS) to one of these regions, Brodmann Area (BA) 17, impairs both imagery and perceptual detection of the same stimuli (Kosslyn et al., 1999). Similarly, activity of primary motor cortex is important for sensori-motor imagery (<u>Pfurtscheller</u> et al., 1997). These results lend weight to the suggestion that regions important for acts of perception and action, can also be important in imagination (Kosslyn et al., 2001).

The role of unimodal cortex in complex internal representations is not limited to states of imagination. For example, contemporary accounts of both memory and emotion emphasize modality specific regions such as those in occipital, auditory, and sensorimotor cortex as playing an important role in generating appropriate mental representations. In both semantic and episodic memory, unimodal regions are assumed to act as ‘spokes’, providing modality specific inputs into ‘hubs’ such as the hippocampus or anterior temporal lobe (Lambon Ralph et al., 2017; Patterson et al., 2007; Moscovitch et al., 2016). These hub regions are closely allied to the default mode network (Davey et al., 2016; de Wael et al., 2018; Murphy et al., 2017; 2018; Raichle et al., 2001; Sormaz et al., 2017;) and play an important role in co-ordinating processing within the spoke regions, allowing representational content to be organised at broader levels. Similarly, accounts of both emotional processing (e.g. Engen, Kanske & Singer, 2016) and interoceptive awareness (Kleckner et al., 2017; Quadt et al., 2018) propose an analogous architecture, with limbic regions such as the amygdala, as well as insular cortex, playing a key role in integrating signals that arise from unimodal cortex. More generally, these functional perspectives are well aligned with topographical views of the cortical hierarchy in which signals arising in peripheral systems, are progressively integrated towards a set of core regions that are equidistant from input and output systems and that help provide a broader, more abstract set of neural operations (Messulam, 1998; Margulies et al., 2016; Paquola et al., 2018).

The current study set out to explore the large-scale functional architecture that underpins acts of imagination. Prior studies have found evidence that certain forms of imagery can recruit brain regions beyond primary systems. For example, Christian et al., (2015) demonstrated that imagining pain from one’s own perspective was linked to greater activity within the anterior insula, and Lucas et al., (2015) demonstrated that the same region responded to both imagined and real touch. Similar patterns of neural recruitment emerge in studies of empathy, which routinely identify that the anterior insula supports representations of pain in another individual (for a review see Bernhardt et al., 2012). Jabbi et al., (2008) documented a role for the anterior insula in experiencing, observing and imagining the experience of disgust. Together these studies provide convergent evidence that at least in certain domains of imagination, regions that are not primarily involved in perception are also important.

The current study aimed to formalise the possibility that aspects of visual imagination depend on a functional network that extends beyond unimodal systems. We combined both task-based and task-free neuroimaging together to allow us to describe the neural architecture that underpins visual imagination. In Experiment 1, we asked a group of participants to imagine familiar everyday images (people, objects and places), as well as viewing pictures of the same stimuli, while neural activity was measured using functional magnetic resonance imaging. This allowed us to establish a set of regions that are important for visual imagination. In Experiment 2, in a large cohort (n = 136), we recorded metrics of brain structure and function and performed a series of cross sectional analyses focused on identifying (i) whether these regions show patterns of intrinsic connectivity at rest and (ii) whether this underlying neural architecture (in both functional and structural aspects) varies with individual differences in the qualities of an individual’s visual imagination. In this way, we hoped to provide converging evidence of the neural architecture that supports visual imagination that takes account of both cortical regions important for visual imagination and population level variance in these attributes. To foreshadow our findings across both studies, we identified regions including the anterior insula and the dorsolateral pre-frontal cortex that together form the ventral attention or saliency network, that play an important role in aspects of visual imagination.

## 2. Methods

### 2.1. Experiment 1 – Task based fMRI

#### 2.1.1. Participants

Forty-one participants were recruited from the University of York. Three participant’s data were excluded due to excessive motion artefacts and an additional two were excluded due to incomplete data sets, leaving thirty-six subjects in the final analysis (22 female; mean age 23.76, range 18-33 years). Participants were native English speakers, right handed and had normal or corrected-to-normal vision. Participants gave written informed consent to take part and were reimbursed for their time. The study was approved by the York Neuroimaging Centre Ethics Committee and Department of Psychology at the University of York. The methods were carried out *in accordance with* the relevant guidelines and regulations outlined in the approved ethics application.

#### 2.1.2. Stimuli

Participants were presented with pictures and written cue words depicting stimuli from three experimental conditions: familiar faces (e.g., David Beckham), places (e.g., Eiffel Tower) and objects (e.g., Microwave). In total there were 45 unique examples; 15 faces, 15 places and 15 objects were shown in both picture and written form. For each unique example (e.g., David Beckham) two different images were presented. Cue words and pictures were presented in blocks for the ‘Seeing’ conditions, whereas only cue words were presented in blocks for the ‘Imagining’ conditions. This yielded six experimental conditions; Seeing Face, Seeing Place, Seeing Object, Imagining Face, Imagining Place, Imagining Object.

#### 2.1.3. Task Procedure

In the scanner there were two runs of 2 blocks (seeing and imagining). Within each run, the ‘imagining’ block was presented first, followed by the ‘seeing’ block. This ensured participants were not simply remembering the exact items from the ‘seeing’ block during imagining. Each block consisted of 45 trials (15 faces, 15 places and 15 objects) presented in a random-order. For the ‘Seeing’ blocks a trial began with a jittered fixation cross (3-4 s), followed by a written cue that identified the upcoming image (e.g., David Beckham) (presented for 1s followed by a jittered fixation cross (0.5-1.5 s), then each of the two images relating to that trial presented for 2 seconds each. On average a trial lasted 10s. A block lasted on average 7 min 30 s. For the ‘Imagining’ block each trial followed the same timing and procedure as ‘Seeing’ blocks, but instead of presenting two images of the target, two unintelligible scrambled images of the target were presented (See Fig 1). Participants were instructed to think about the item depicted in the written cue during the presentation of either images (seeing blocks) or scrambled image (imagining blocks). In addition, participants were instructed to press a button whenever a red fixation-cross was presented to ensure they were paying attention. This occurred randomly on 20% of trials.

**Figure 1.**
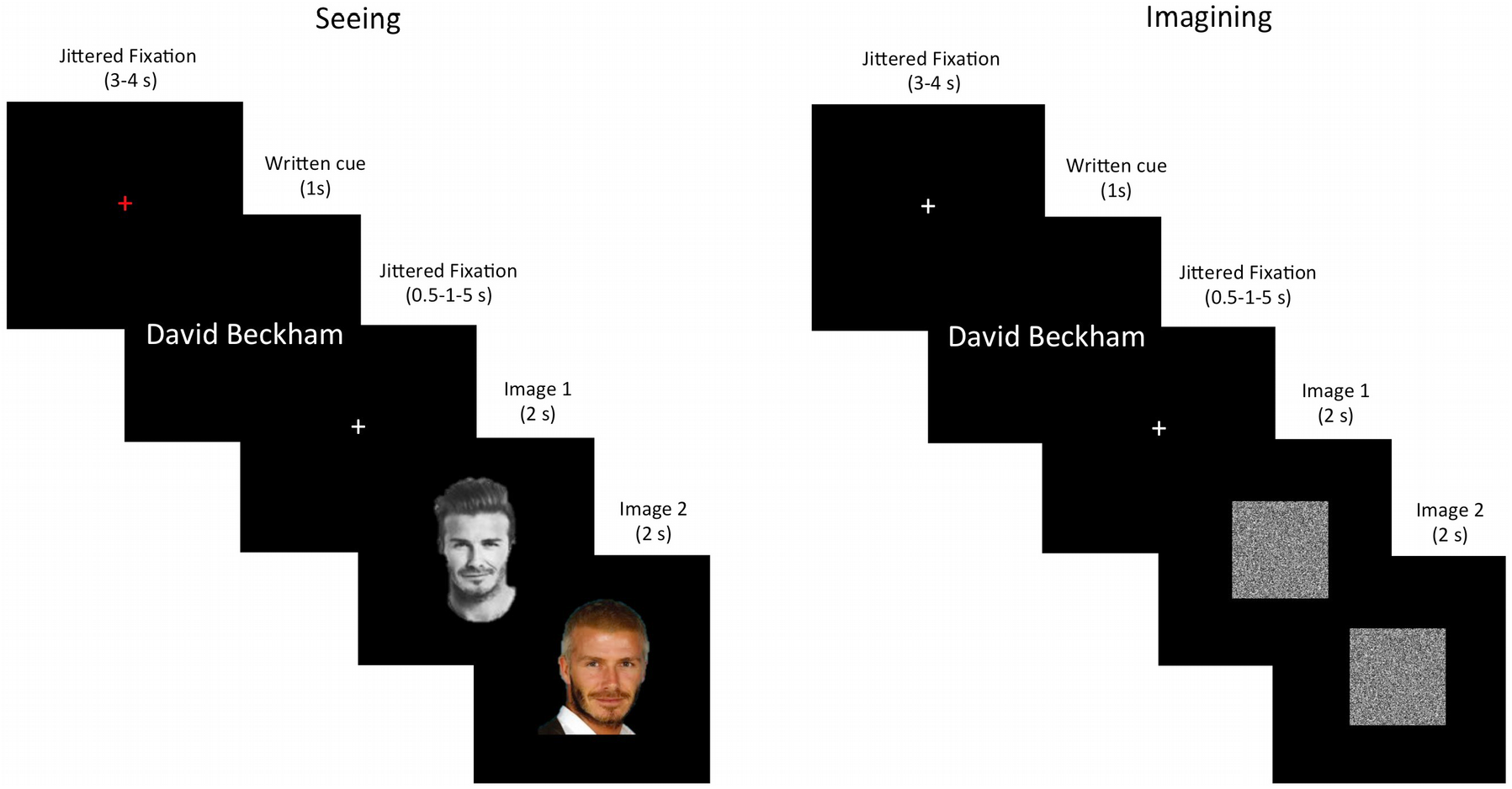
Experimental Design. Example of one face trial for both ‘seeing’ and ‘imagining’ blocks. Each trial lasted on average 10 seconds. For each block, 15 face trials, 15 scene trials and 15 object trials were presented in a random order (n=45 in total). A block lasted on average 7 min 30s. For both trials participants were instructed to think about the item depicted in the written cue during the presentation of either images (seeing blocks) or scrambled image (imagining blocks). In addition, participants were instructed to press a button whenever a red fixation-cross was presented to ensure they were paying attention. This occurred randomly on 20% of trials.

After being scanned, participants completed a questionnaire rating their familiarity, personal relevance and imageability for each of the 45 items (15 faces, 15, places, 15 objects). These ratings were done on a 7-point likert scale.

#### 2.1.4. Acquisition

Data were acquired using a GE 3T HD Excite MRI scanner at the York Neuroimaging Centre, University of York. A Magnex head-dedicated gradient insert coil was used in conjunction with a birdcage, radio-frequency coil tuned to 127.4 MHz. A gradient-echo EPI sequence was used to collect data from 60 bottom-up axial slices (TR = 3 s, TE = minimum full, matrix size = 64 x 64, flip angle = 90°, voxel size = 3 x 3 x 3 mm). Functional images were co-registered onto a T1-weighted anatomical image from each participant (TR = 7.8 s, TE = 3 ms, FOV = 290×290 mm, matrix size = 256 x 256 mm, voxel size = 1 x 1 x 1 mm) using a linear registration (FLIRT, FSL). A FLAIR scan with the same orientation as the functional scans was collected to improve co-registration between scans.

#### 2.1.5. Pre-processing

Task based imaging data were pre-processed using the FSL toolbox (http://www.fmrib.ox.ac.uk/fsl). Images were skull stripped using a brain extraction tool (BET, Smith, 2002) to remove non-brain tissue form the image. The first 3 volumes (6 s) of each scan were removed to minimize the effects of magnetic saturation, and slice-timing correction was applied. Motion correction (MCFLIRT, Jenkinson et al., 2002) was followed by temporal high-pass filtering (cutoff = 0.01Hz). Individual participant data were first registered to their high-resolution T1-anatomical image using the linear registration tool FLIRT, and then into a standard space (Montreal Neurological Institute (MNI152); this process included tri-linear interpolation of voxel sizes to 2×2×2x mm. Finally, we applied a Gaussian smoothing kernel with a full width half maximum of 6 mm.

#### 2.1.6. Univariate analysis

The trial onset was taken from the presentation of the first image for each trial, and the duration was set at 4s. Response to the red fixation cross was also modelled as a regressor of no interest. The response to each trial was contrasted against rest. Box-car regressors for each condition (Seeing Faces, Seeing Places, Seeing Objects, Imagining Faces, Imagining Places, Imagining Objects), for each run, in the general linear model were convolved with a double gamma hemodynamic response function (FEAT, FSL). Regressors of no interest were also included to account for head motion within scans. A fixed effect design (FLAME, FSL) was then conducted to average across the two runs, within each individual. Finally, individual participant data were entered into a higher-level group analysis using a mixed effects design (FLAME, FSL) whole-brain analysis.

### 2.2. Experiment 2 - Resting state fMRI

#### 2.2.1. Participants

This analysis was performed on a separate cohort of 136 healthy participants at York Neuroimaging Centre (51 male; mean age 20.26, range 18-31 years). Subjects completed a 9-minute resting-state rfMRI (rs-fMRI) scan during which they were asked to rest in the scanner with their eyes open. Using these data we examined the rs-fMRI connectivity of regions that were informative to seeing images (seeing > imagining) and imagining (imagining > seeing) to investigate whether these regions fell within similar or distinct networks. These participants also completed the Evaluation of Mental Imagery questionnaire which evaluated four domains of mental imagery (Christian, Miles, Parkinson, & Macrae (2013): (1) time orientation of mental imagery (past-future), (2) emotional valence of mental imagery (negative-positive), (3) vividness of mental imagery (not vivid-extremely vivid) and (4) visual perspective of mental imagery (3^rd^ person - 1^st^ person perspective).

#### 2.2.2. Acquisition

As with the functional experiment, a Magnex head-dedicated gradient insert coil was used in conjunction with a birdcage, radio-frequency coil tuned to 127.4 MHz. A gradient-echo EPI sequence was used to collect data from 60 bottom-up axial slices (TR = 3 s, TE = minimum full, FOV = 192×192 mm, matrix size = 64×64, voxel size = 3×3×3 mm, flip angle = 90°). A minimum full TE was selected to optimise image quality (as opposed to selecting a value less than minimum full which, for instance, would be beneficial for obtaining more slices per TR). Functional images were co-registered onto a T1-weighted anatomical image from each participant (TR = 7.8 s, TE = 3 ms, FOV = 290×290 mm, matrix size = 256×256 mm, voxel size = 1×1×1 mm) using a linear registration (FLIRT, FSL).

#### 2.2.3. Pre-processing

Resting-state data pre-processing and statistical analyses were carried out using the SPM software package (Version 12.0), based on the MATLAB platform (Version 15a). For preprocessing, functional volumes were slice-time and motion-corrected, co-registered to the high-resolution structural image, spatially normalised to the Montreal Neurological Institute (MNI) space using the unified-segmentation algorithm (Ashburner & Friston, 2005), and smoothed with an 8 mm FWHM Gaussian kernel. With the goal of ensuring that motion and other artefacts did not confound our data, we first employed an extensive motion-correction procedure and denoising steps, comparable to those reported in the literature (Ciric et al., 2017). In addition to the removal of six realignment parameters and their second-order derivatives using the general linear model (GLM) (Friston et al., 1996), a linear detrending term was applied as well as the CompCor method that removed five principle components of the signal from white matter and cerebrospinal fluid (Behzadi et al., 2007). Moreover, the volumes affected by motion were identified and scrubbed based on the conservative settings of motion greater than 0.5 mm and global signal changes larger than z = 3. A total of fifteen participants, who had more than 15% of their data affected by motion was excluded from the analysis (Power et al., 2014). Though recent reports suggest the ability of global signal regression to account for head motion, it is also known to introduce spurious anticorrelations, and thus was not utilised in our analysis (Saad et al., 2012). Finally, a band-pass filter between 0.009 Hz and 0.08 Hz was employed in order to focus on low frequency fluctuations (Fox et al., 2005).

Following this procedure, two seed regions were generated from the univariate map of seeing>imagining and imagining>seeing (see Figure 4; Table 1). This yielded two peak clusters centred around left lateral occipital cortex for seeing>imagining [−44 −76 −12] and occipital pole for imagining>seeing [0 −98 14]. The Conn functional connectivity toolbox (Version 15.h) (Whitfield-Gabrieli & Neston-Castanon, 2012) was used to perform seed-based functional connectivity analyses for each subject using the average signal from the spheres placed on the MNI coordinates for the two regions of interest (ROIs) described above. In addition, each of the four Z-scored results from the mental imagery questionnaire were used as independent behavioural regressors to interorgate whether response modulated resting-state connectivity of each of our seed regions. All reported clusters were corrected for multiple comparisons using the Family Wise Error (FWE) detection technique at the .05 level of significance (uncorrected at the voxel-level, .001 level of significance).

### 2.3. Meta analytic decoding

To allow quantitative inferences to be drawn on the functional neural activity identified through our seed based correlational analyses we performed an automated meta-analysis using NeuroSynth (used 2018; http://neurosynth.org/decode; <u>Yarkoni et al., 2011</u>). Using an association test, this software computed the spatial correlation between each RS mask and every other meta-analytic map (n=11406) for each term/concept stored in the database (e.g., vision, imagery, memory, decision making). The 10 meta-analytic maps exhibiting the highest positive correlation and for each sub-system mask were extracted, and the term corresponding to each of these meta-analyses is shown in <u>Fig</u>. 2. The font size reflects the size of the correlation. This allows us to quantify the most likely reverse inferences that would be drawn from these functional maps by the larger neuroimaging community.

**Figure 2.**
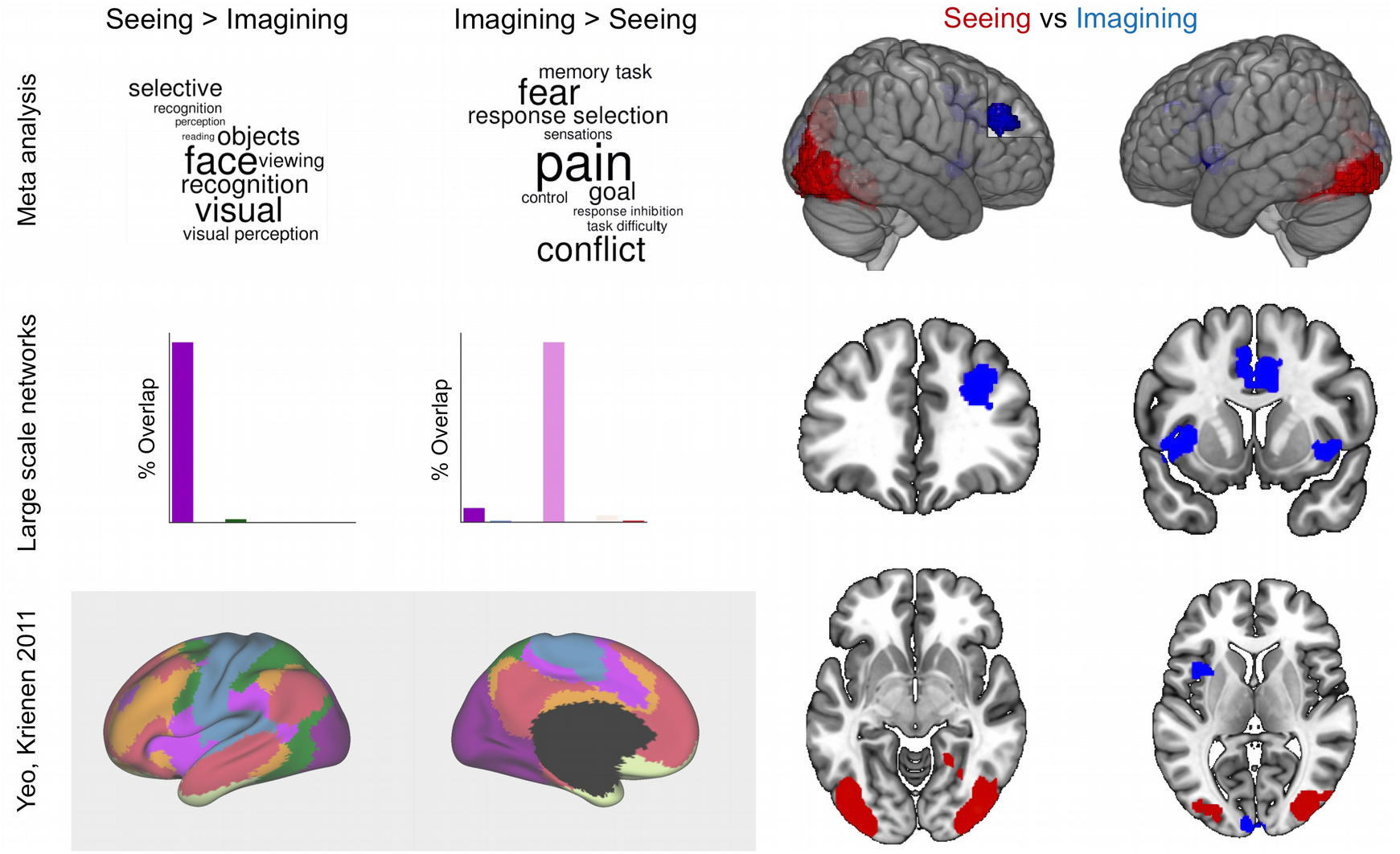
Experiment 1. Regions showing stronger activation during seeing (red) and imagining (blue) familiar people, places and objects. Bar charts show the percentage of voxels that fall within different large scale networks (see Yeo, Krienen et al., 2011). Word clouds describe the results of a meta-analytic decoding of the spatial maps using Neurosynth. Spatial maps were thresholded at Z = 3.1, and family wise error is controlled at p<.05.

### 2.4. Cortical thickness analysis

#### 2.4.1. Pre-processing

Cortical thickness is modelled from the T1-weighted anatomical image acquired with the resting state data using models generated by FreeSurfer (5.3.0; https://surfer.nmr.mgh.harvard.edu/), a tool widely validated on its accuracy of the thickness measures (Fischl, 2012). The following preprocessing steps were included (for details, please refer to Dale et al., 1999; Fischl et al., 1999): intensity normalization, removal of nonbrain tissue, tissue classification, and surface extraction. The extracted surfaces was aligned with curvature of an average spherical representation, fsaverage5, to improve the correspondence of measurement locations among subjects. Each of these surfaces from individual subjects was also visually inspected and, if necessary, manually corrected. Finally, cortical thickness was calculated based on the closest distance between the grey/white boundary and pial surface at each vertex across the entire cortex. A surface-based smoothing with a 20 mm full-width-at-half-maximum (FWHM) Gaussian kernel was applied to reduce measurement noise without forgoing the capacity for anatomical localization (Lerch and Evans, 2005).

#### 2.4.2. Seed-based analysis

Five ROI defined in volume space, taken from from the univariate map of imagining>seeing (see Figure 3; Table 1 ??), was submitted to FreeSurfer to extract the corresponding cortical thickness values. This was done for each ROI, that is, we created an fsaverage5-ROI surface overlay by mapping the volumetric ROI to an anatomical volume registered to the freesurfer template subject, fsaverage5, and then extracted the mean cortical thickness value by mapping the fsaverage5-ROI surface overlay onto the thickness data of individual subject. Mean cortical thickness for all five ROIs were interrogated for possible relationship with the four domains of mental imagery. To perform this analysis we first applied principal components analysis to the matrix of cortical thickness for each participant. This revealed two components with eigen variates greater than 1 (see Results). These two components were included as between participant regressors in a mixed ANOVA with the scores on each questionnaire item as dependent variables.

**Figure 3.**
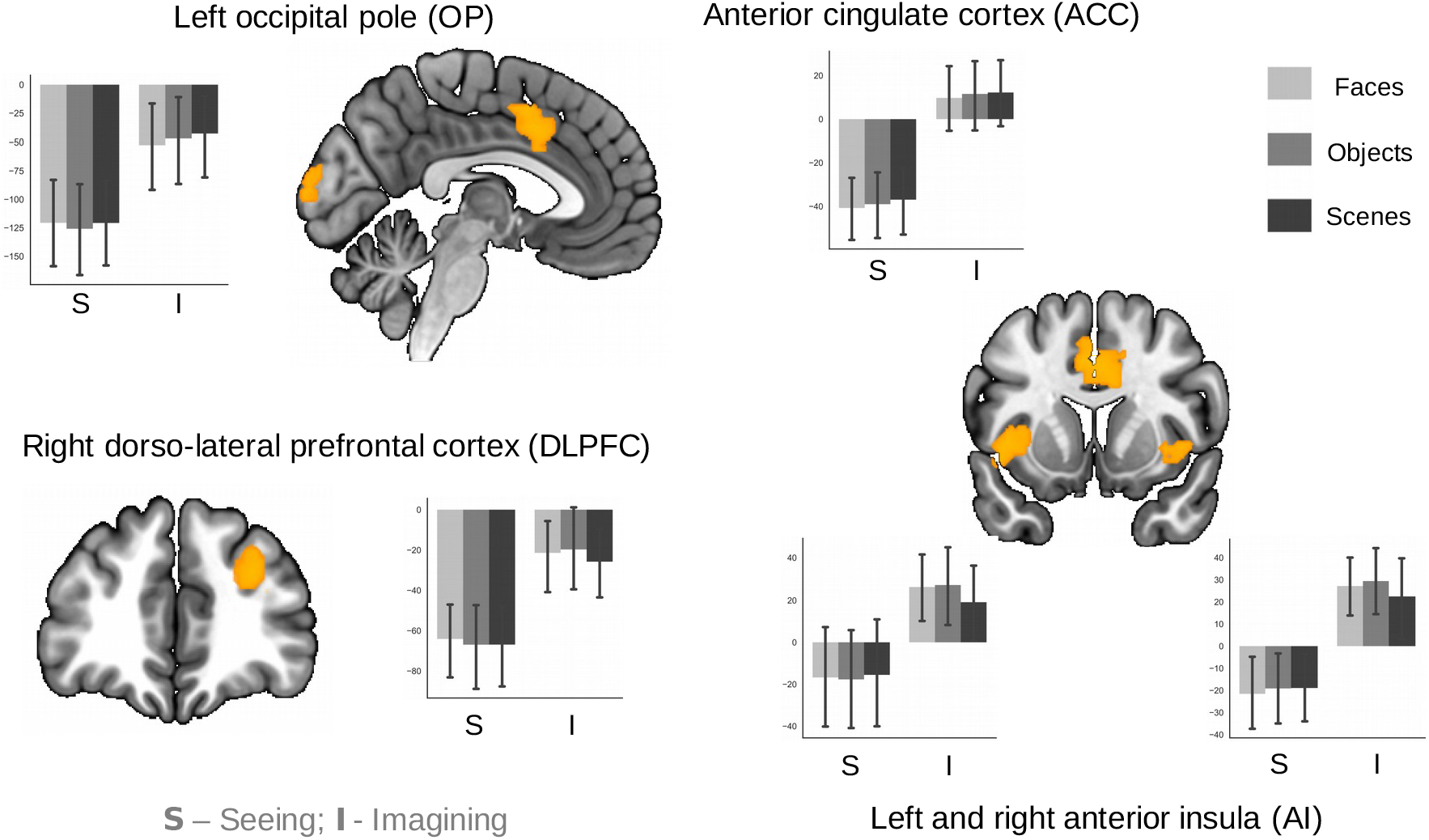
Experiment 1. Regions showing patterns of reduced deactivation, as well as heightened activity during imagination relative to seeing.

## 3. Results

Our first analysis used task-based data recorded in Experiment 1 to compare patterns of neural activity observed during imagining or seeing the familiar people, places and objects. Contrasting imagination over seeing yielded five clusters of neural activity. These included the left occipital cortex (BA 17), the left and right anterior insula (BA 13), the anterior cingulate cortex (BA 32) and the right dorsolateral prefrontal cortex (BA 9/46). The reverse comparison (seeing greater than imagination) yielded a bilateral cluster of activity within lateral visual cortex (BA 19/37). These results are presented in Figure 2. We also performed a meta-analytic decoding of these regions using Neurosynth (Yarkoni et al., 2011) to help evaluate their most likely functional associations. Regions active when viewing vs imagining, yielded terms related to visual perception including “visual”, “recognition”, and also content specific terms that partly matched what our participants viewed - “faces” and “objects”. In contrast, regions associated with imagination had a broader functional profile including affective terms such as “pain” and “fear”, as well as domain general processes such as “response selection”, “conflict” and “goal”. Finally, a comparison with a popular set of large-scale networks (Yeo et al., 2011) indicated that regions sensitive to seeing falls largely within the visual network, while the regions sensitive to imagination fall within a small region of visual cortex, with the majority falling within the ventral attention or salience network.

We examined the percent signal change in each region with a stronger response during imagination than seeing (see Figure 3). Neural response within the occipital pole and the dorsolateral prefrontal cortex showed patterns of reduced deactivations during imagination relative to seeing. In contrast, the anterior cingulate cortex, as well as the bilateral anterior insula tended to show increases in activity above baseline during imagination. It can also be seen in all regions that neural responses were similar regardless of the type of content that was imagined (faces, places and objects).

Having identified a set of regions showing patterns of increased activity during acts of imagination, we next used the resting-state data recorded in Experiment 2 to explore their intrinsic architecture. We wanted to evaluate whether the regions of unimodal cortex involved in imagination (i.e. BA 17) reflect a region of visual cortex that show greater functional overlap with the more distant regions we also observed as active during imagination (i.e. the elements of the ventral attention network). To examine this question we contrasted the connectivity of areas of occipital pole, showing greater activity during imagination, with lateral regions of occipital cortex (i.e. BA 18 and 37) showing higher neural responses while observing the same stimulus (see Methods). These spatial maps highlight regions more connected to portions of medial visual cortex important for imagination, as well as those connected to lateral visual regions that show a greater response to visual input (see left hand panel of Figure 4). Notably, our medial region of visual cortex was more connected to the cingulate cortex, regions of angular gyrus and dorsolateral prefrontal cortex. In contrast, lateral visual cortex was more connected to sensorimotor cortex, ventral prefrontal, as well as regions of dorsal parietal cortices.

**Figure 4.**
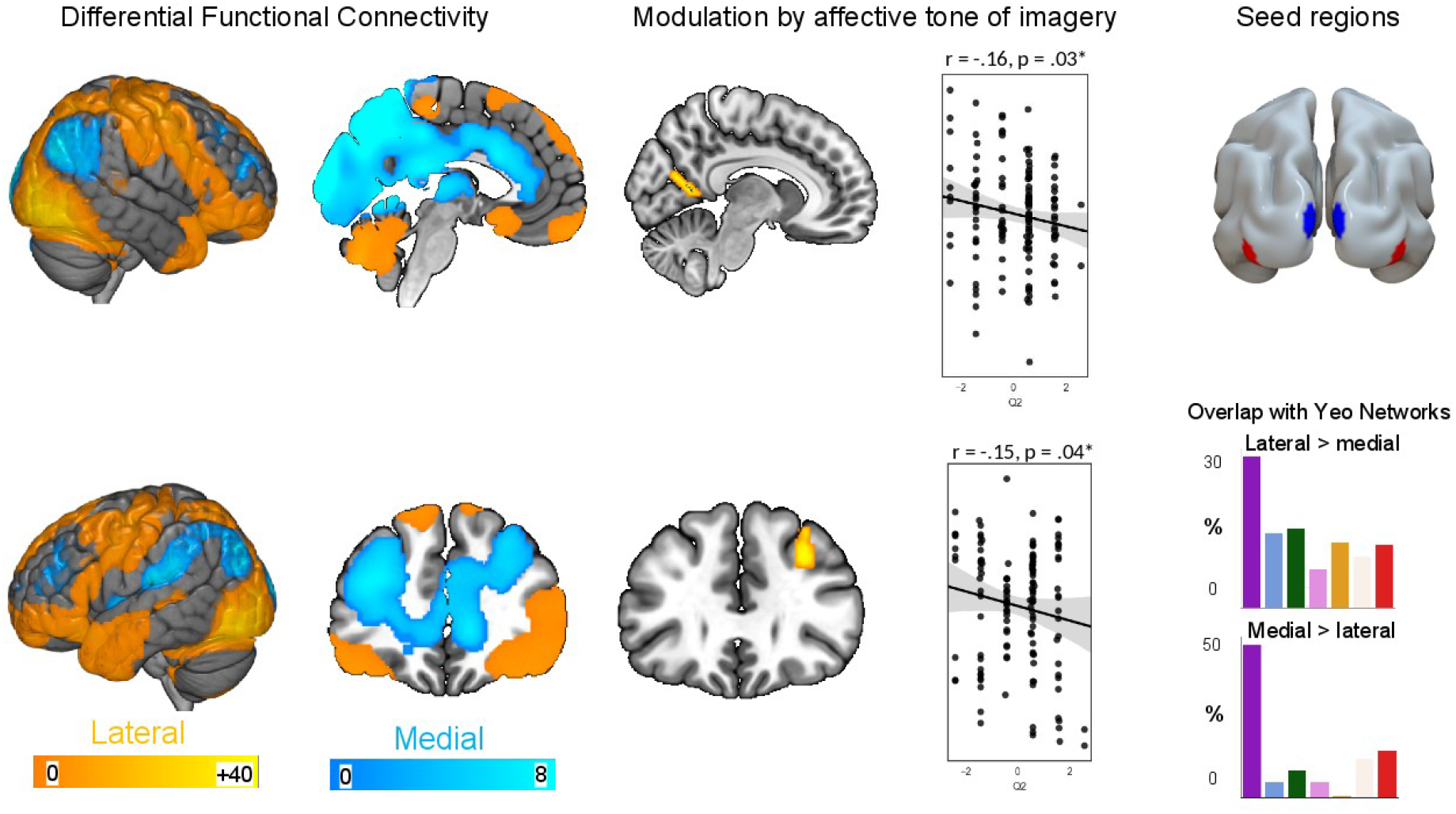
Experiment 2. *Left hand panel*. Regions showing differential connectivity with medial visual regions showing higher activity during imagination and lateral regions of visual cortex that are more active during seeing. *Middle panel*. Regions showing regions modulation of this pattern of differential connectivity based on the emotionality of individual self-reported patterns of visual imagery. The scatter plots show each individual. *Right hand panel*. The medial and lateral seed regions used in this analysis.

We hypothesized that this pattern of intrinsic architecture has a functional relationship to individual variation in the nature of imagery. To test this possibility we examined whether the intrinsic architecture generated in the last step of our analyses was related to individual differences in how these experiences unfold. We repeated the differential connectivity analysis described above, including each individual’s score on a validated measure of visual imagery as an additional variable of interest (Christian et al., 2013). We found that the differential connectivity of the two regions of medial visual cortex was correlated with the affective style of an individuals’ visual imagery. The results of this analysis are presented in the middle panel of Figure 4 where it can be seen that this was localised to a region of retrosplenial cortex, as well as region of dorsolateral prefrontal cortex. Specifically, these results indicated individuals for whom visual imagery tended to be negative in tone, showed stronger coupling between these regions and medial visual cortex (see scatterplot).

Next we examined the relationship between the nature of ongoing imagination and individual differences in cortical thickness. We extracted the cortical thickness within the regions identified as important in generating images in imagination and then examined their association with the descriptions of imagery in daily life. For the purposes of analysis we first decomposed the cortical thickness data revealing two significant components. These data were analyzed using a repeated measures analysis with each of the questionnaire scores describing the different types of imagery as dependent variables and each loadings on the principal components as variables of interest, as well as age and gender as nuisance variables. This found a significant interaction between the first component, and the individuals likelihood of their imagery adopting a third-person perspective, showing greater cortical thickness on average across the set of regions activated in Experiment 1 [F (1, 122) = 4.6, p<.05] (see Fig. 5).

**Figure 5.**
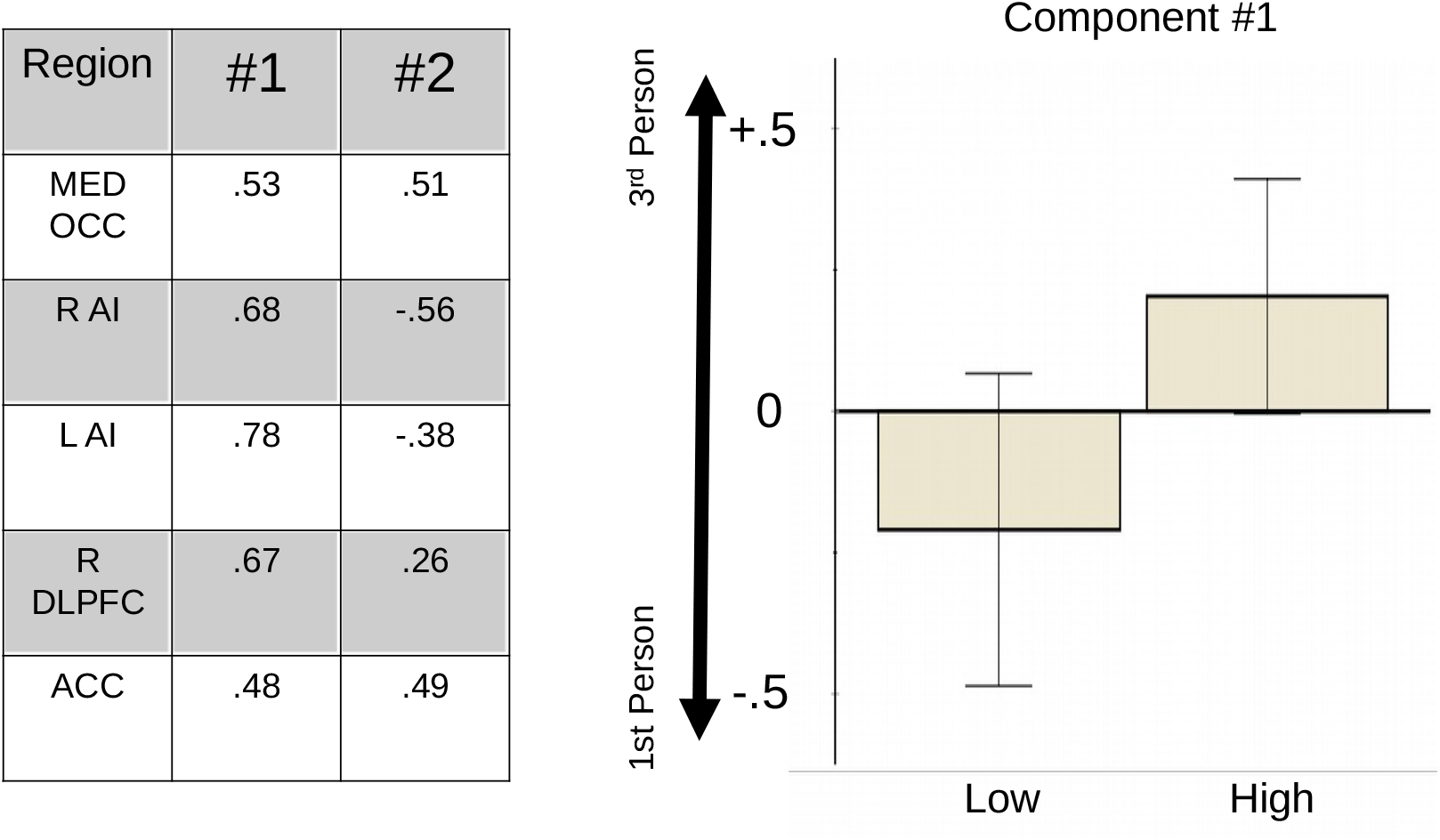
Experiment 2. Associations with cortical thickness and the visual perspective adopted during visual imagery. A decomposition of the cortical thickness in the regions linked to visual imagery in Experiment 1 revealed two components with eigen variates greater than one. Individual’s loading higher on the first component were more likely to report imagery, which often took a third-person perspective. The error bars are the 95% confidence intervals.

In our final analysis, we consider the relationship between the results of our two experiments. First, we examine whether the relationship between the distributed pattern of regions showing evidence of greater activity during imagination from Experiment 1, with the spatial distribution of the connectivity of the medial region observed in Experiment 2. It can be seen in the right-hand panel of Figure 6 that the spatial maps overlapped within three of the four regions showing stronger activity during imagination (anterior cingulate cortex, dorsolateral prefrontal cortex and left anterior insula). This analysis demonstrates that despite the distributed nature of regions recruited during visual imagination, analysis of their intrinsic architecture suggest that they are all closely coupled at rest to the region of primary visual cortex showing similar activity during acts of imagination. Next, we explored whether the patterns of increased activity seen during Experiment 1 were linked to the regions associated with greater connectivity for individuals with more negatively toned visual imagery. These spatial maps overlapped in the same region of dorsolateral prefrontal cortex as was observed in our prior analyses (see middle and right-hand columns in Figure 6). A formal conjunction of these maps (using FSL easythreshconj command) revealed a significant conjunction (z=3.1, p < .01) as seen in the middle panel of Figure 6. These results show that a stronger pattern of intrinsic organisation between regions identified as having greater neural activity during imagination, is related to the emotional qualities of visual imagery, thus underlining the broader architecture identified in our study has implications for the way that imagery unfolds in the psychological domain.

**Figure 6.**
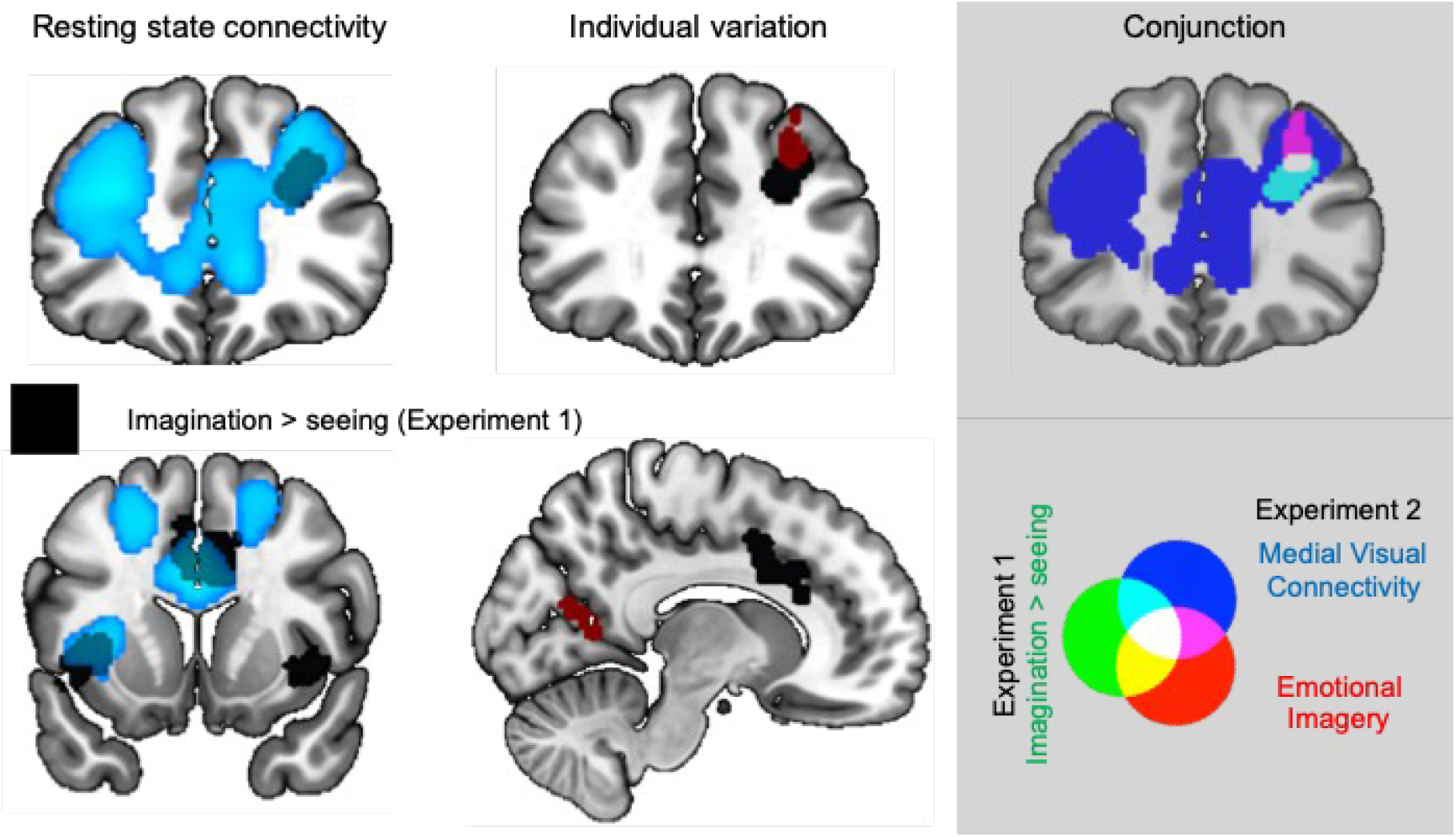
Experiment 1 and 2. Converging evidence from for a role of right dorsolateral prefrontal cortex in visual imagination. *Left hand column*. Regions of visual cortex showing greater neural activity during visual imagination relative to seeing show heightened connectivity with other areas of cortex active at rest. *Middle column*. A region of dorsolateral prefrontal cortex showing heightened connectivity for individuals whose imagery had a more negative affective tone overlapped with a region showing heightened activity during imagination relative to seeing. *Right hand column*. A region of right dorsolateral prefrontal cortex was common to both analyses. All spatial maps were thresholded with a cluster forming threshold of Z = 3.1 and corrected for family wise error at p <.05.

## 4. General discussion

In two experiments we explored whether the neural substrates underlying visual imagination can be understood from the perspective of a component process architecture. In Experiment 1 we replicated prior findings that BA17 is involved in visual imagery, and found that activity also increased within core regions of the ventral attention network (anterior insula, dorsolateral prefrontal cortex and the pre-supplementary motor area). In Experiment Two, task-free fMRI demonstrated that the network of regions activated during imagination were functionally coupled at rest, and that the strength of the coupling between BA17 and the dorsolateral prefrontal cortex was related to the emotional qualities of an individuals visual imagery. Finally, we found that cortical thickness within this set of regions was greater for individuals whose visual imagination often spanned a third-person perspective. Together, these findings highlight that the process through which we visualise information in imagination depends on a functional network involving both primary sensory cortex, and aspects of the cortex in particular within the ventral attention network.

Our study highlights that visual imagination, like other aspects of higher order cognition, such as semantic and episodic memory, relies on a distributed network that spans both a primary unimodal system (in this case BA17) as well as regions of heteromodal cortex within the ventral attention network. A role for the ventral attention network in visual imagination is interesting given that the other canonical attentional system, the dorsal attention network, is closely tied to visual processing (see for example Braga et al., 2016; Murphy et al., 2018). Regions within the dorsal attention network are important in the maintenance of attention on a visual field (Kastner et al., 1999) and key nodes of this network contain retinotopic maps (Moore et al., 2003, Ruff et al., 2008). Unlike the dorsal attention network, the ventral attention network plays a more general role enhancing patterns of cognition that are specific to a given goal context. For example, the ventral attention network exhibits a pattern of sustained activity during task sets (Dosenbach et al., 2006). Importantly, we have recently found a similar region of left dorsolateral prefrontal cortex that plays an important role in prioritizing patterns of self-generated thoughts when task demands are low (Turnbull et al., 2018, 2019). The fact that our study highlights the ventral rather than dorsal attention network, as important in visual imagination, adds to a growing body of evidence that the ventral attention system plays a role in prioritizing internal states as well as external task processing.

The results of Experiment 2 provide additional support for the role of the ventral attention network during visual imagination, particularly when these experiences are highly emotional. Our functional analysis shows that variation in the emotional tone of visual imagery was associated with stronger connectivity between regions of medial visual cortex and regions of the dorsa-lateral cortex. Studies indicate that the generation of affectively laden images is a hallmark of both normal and abnormal patterns of experience. For example, accounts of patterns of ongoing thought often studied under the rubric of mind-wandering highlight that the content of these experiences often relate to an individual’s current concerns – “a latent, time-binding brain process (a current concern) that sensitizes the individual to notice, recall, think about, dream about, and act on cues associated with the goal pursuit” (Klinger and Cox, 2004, page 3). In normal individuals, the notion of current concerns helps explain why much spontaneous mentation is related to the self and other individuals (e.g. Stawarczyk et al., 2013). In addition, studies of the tendency to engage in spontaneous mentation is enhanced by negative moods (e.g. Smallwood et al., 2009) and chronic states of negative affect are associated with exacerbated levels of mind-wandering (Smallwood et al.,, 2007). Together our data suggests that one role that the ventral attention network plays in visual imagination is by helping to actively prioritise information that is relevant to the individual, and that this process is strengthened by particularly salient aspects of imagination. This view is consistent with evidence that the ventral attention network helps prioritise salient stimuli across multiple domains including pain (Atlas et al., 2010), music (Sridharan, et al., 2007) and information important for social interactions (for a review, see Decety and Lamm, 2007). Notably, an emerging literature suggests that neurostimulation of regions of pre-frontal cortex is helpful in the amelioration of depressive symptoms (e.g. Mak et al., 2017). Since visual images with a negative tone are common in depression (for a review, see Holmes et al., 2016), it is possible that the reason why stimulation of dorsolateral prefrontal cortex can alleviate depression is because it helps the individuals exert control over emotional qualities of their self-generated experiences.

In addition, our study shows that cortical thickness within regions of the ventral attention network is greater for individuals who tend to experience patterns of imagery from a third-person perspective (rather than from a first-person view). It is possible that this pattern also reflects a role of salience since in conditions such as social phobia, individuals often experience intrusive thoughts about how other individuals view them (e.g. Spurr and Stopa, 2003). Alternatively, it could reflect the additional processing resources involved in imaging information from the perspective of other individuals. For example, it has been argued that a third-person perspective may require additional propositional information that is not necessary for imagery based on an egocentric view point (Libby and Eibach, 2011; Libby et al., 2014). Future work could compare these different perspectives by examining whether the complexity of imagination from a third person perspective, or its relevance to social processes that reflect this association.

At a more mechanistic level, our study suggests the ventral attention network plays an important role in visual imagination and we close by considering the neurocognitive mechanisms that may underlie this process. One region identified in both experiments - the right dorsolateral prefrontal cortex - is known to be important in prioritising both internal and external sources of information in a goal directed manner (Petrides, 2005; Jiang 2018). Furthermore, it is hypothesised that the capacity of dorsolateral prefrontal cortex to promote goal relevant information from both internal and external domains, provides a zone of contextual control that helps sustain goal relevant processes in a flexible manner (Badre and Nee, 2018). We speculate that humans exploit the ability of the dorsolateral prefrontal cortex to co-ordinate multiple sources of internal and external information to allow the visualisation of people, places and objects that are not present in the here and now. A second region of the ventral attention network, the right anterior insula, was also highlighted as important in visual imagination by both experiments. This region is an important hub in integrating interoceptive information with signals from elsewhere in the cortex (Craig, 2009). Consistent with our analyses, studies have highlighted the anterior insula as important for adopting another person’s perspective than your own in both assessing another’s pain (Singer et al., 2004; Jackson et al.,, 2005) as well as when deciding about charitable donations (Tusche et al., 2016).

## Acknowledgements

Thanks to York Neuroimaging Centre for technical support.

## Funding

This project was supported by European Research Council Consolidator awarded to JS (WANDERINGMINDS – 646927).

## Author contributions

C.M., M.S and J.S. conceived of the ideas for analysis. Computations were performed by C.M. with assistance from N. S. P. H., (structural data) B.B., C. N. M., M.S., and D. V. (functional data); The manuscript was written by C.M. and J.S. with contributions from B.B., C.N.M., E.J., and D.V.

## Competing interests

Authors declare no competing interests.

## Data availability statement

Raw Z-maps from the task-based analysis are available on Neurovault in a collection with the title of this article, along with the significant components from the resting state and cortical thickness analyses. All summary data and materials used in the analysis are available on request. Raw fMRI data is restricted in accordance with ERC and EU regulations.

## Code availability statement

All code used in the production of this manuscript is available on request.

